# A cattle-derived human H5N1 isolate suppresses innate immunity despite efficient replication in human respiratory organoids

**DOI:** 10.1101/2025.11.02.684669

**Authors:** Shintaro Shichinohe, Hikaru Sugimoto, Masako Yamasaki, Rina Hashimoto, Tatsuru Morita, Daiki Kobayashi, Takahiro Hiono, Daisuke Motooka, Makoto Shimooka, Norikazu Isoda, Mai Thi Quynh Le, Ayato Takada, Yoshihiro Sakoda, Eiryo Kawakami, Kazuo Takayama, Tokiko Watanabe

**Author notes:** These authors contributed equally (Shintaro Shichinohe, Hikaru Sugimoto). Correspondence (E.K.), (K.T.) (T.W.).

## Abstract

The H5N1 high pathogenicity avian influenza virus (HPAIV) of clade 2.3.4.4b, which spreads globally via wild birds, has become a major public health concern because it can infect a variety of mammals, including humans. In 2024, infection of dairy cattle with H5N1 HPAIV clade 2.3.4.4b was confirmed in the United States, and subsequent human cases were reported. Although these viruses are highly pathogenic in animal models, human infections have generally been mild, revealing a striking discrepancy. Here, we characterized the cattle-derived human H5N1 isolate A/Texas/37/2024 (TX37-H5N1) using three-dimensional human respiratory organoids derived from induced pluripotent stem (iPS) cells. Despite efficient replication, TX37-H5N1 induced minimal interferon and inflammatory cytokine responses. Bulk and single-cell RNA sequencing revealed reduced STAT1-mediated transcriptional activity in TX37-H5N1-infected organoids compared to the historic H5N1 human isolate A/Vietnam/1203/2004. These findings suggest that TX37-H5N1 fails to trigger the strong innate responses, including robust cytokine production, that are typically associated with severe H5N1 disease and are thought to contribute to cytokine storm-medicated pathogenesis. This attenuated response may help explain the discrepancy between the high pathogenicity of TX37-H5N1 in animal models and its mild clinical presentation in humans. While zoonotic influenza risk is often assessed using cell lines or animal models, our study highlights the value of using human respiratory organoids to evaluate human-specific virus-host interactions. This platform provides a complementary tool for assessing the risk of emerging avian influenza viruses.

**Highlights:**

- Human respiratory organoids were used to model zoonotic B3.13 H5N1 infections.
- A cattle-derived human isolate, TX37-H5N1, replicated more efficiently than historical VN1203-H5N1.
- TX37-H5N1 suppressed STAT–IRF-mediated innate immune responses.
- TX37-H5N1 was sensitive to baloxavir and oseltamivir but less sensitive to favipiravir.
- Human respiratory organoids offer a complementary platform for zoonotic influenza risk assessment.

**In brief:** Our study used human iPSC-derived respiratory organoids to investigate the mild clinical presentation of zoonotic B3.13 H5N1 viruses. We found that TX37-H5N1 replicates efficiently but suppresses innate immune responses, providing mechanistic insight into species-specific pathogenesis and highlighting the utility of human organoids for zoonotic risk assessment.

## Introduction

Avian influenza viruses are zoonotic pathogens with pandemic potential that circulate in wild birds and occasionally spill over into mammals, including humans. Among them, H5N1 highly pathogenic avian influenza viruses (HPAIVs) have historically caused severe disease in humans, with sporadic infections and high case fatality rates since their emergence in 1997. Modern H5N1 HPAIVs, belonging to the Goose/Guangdong (Gs/GD) lineage, have diversified into multiple genetic clades, with clade 2.3.4.4b currently dominating global outbreaks through migratory bird transmission and causing unprecedented infections across a diverse array of mammalian hosts^1^. Notably, in 2024, clade 2.3.4.4b H5N1 HPAIV infections were reported for the first time in dairy cattle in the United States^2^, followed by zoonotic human infections^3^. These viruses belong to the genotype B3.13 viruses, which emerged through reassortment with North American wild bird strains^1^. Interestingly, human infections with B3.13 viruses have been largely limited to conjunctivitis or mild respiratory symptoms^3,4^, in stark contrast to the high case fatality rate (approximately 48%) of earlier H5N1 HPAIVs^5^.

H5N1 HPAIVs have caused severe disease in humans and animal models (e.g., mice, ferrets, and non-human primates), with high case fatality rates associated with hyperinflammatory responses and acute respiratory distress syndrome (ARDS). Studies in mice have shown that H5N1 infection induces strong activation of STAT1 and IRF signaling pathways, which are implicated in the excessive cytokine responses characteristic of severe disease^6^. Recent studies have demonstrated that B3.13 viruses retain high virulence in animal models, including mice, ferrets^7,8^, and non-human primates^9^, yet cause only mild symptoms in humans. This striking discrepancy raises fundamental questions about the molecular determinants of pathogenesis and whether B3.13 viruses fail to induce the cytokine storms typically associated with severe H5N1 disease^10^.

Traditional animal models can be limited in that they capture species-specific host responses. Recent advances in human induced pluripotent stem cell (iPSC)-derived respiratory organoids offer a physiologically relevant 3D model system that recapitulates the multicellular architecture and immune responsiveness of human airway tissues. Importantly, the organoid system we used in the present study incorporates not only epithelial cells but also non-epithelial populations such as pericytes, which contribute to the complex microenvironment and host responses. This organoid platform has been successfully applied to study host-pathogen interactions in SARS-CoV-2 and respiratory syncytial virus (RSV), underscoring its utility in dissecting human-specific viral dynamics and immune responses.

In this study, we sought to elucidate the molecular basis of the mild clinical outcomes associated with B3.13 H5N1 infections in humans. We compared a cattle-derived human isolate, A/Texas/37/2024 (referred as TX37-H5N1), with an historic highly pathogenic human H5N1 isolate, A/Vietnam/1203/2004 (referred as VN1203-H5N1), to investigate differences in viral replication, cellular tropism, and host immune responses. By employing iPSC-derived respiratory organoids that recapitulate the multicellular complexity of human respiratory tissues, we demonstrate the physiological relevance of this platform to dissect human-specific antiviral responses. Our findings offer new insights into the host-pathogen interactions that may explain the apparent reduced clinical severity of recent zoonotic infections. Furthermore, this study establishes human organoid-based approaches as valuable complementary tools for assessing the pandemic risk of emerging avian influenza viruses, providing key molecular insights into human-specific antiviral responses.

## Results

### TX37-H5N1 HA exhibits receptor-binding properties typical of avian influenza viruses

Influenza virus tropism and pathogenesis are largely determined by the receptor-binding specificity of viral hemagglutinin (HA), which recognizes sialic acid-containing glycans on host cell surfaces. To evaluate receptor-binding specificity, we generated recombinant HAs from the cattle-derived H5N1 human isolate TX37-H5N1 and an historic H5N1 HPAIV isolated from a fatal human case [A/Vietnam/1203/2004 (VN1203-H5N1)]. Using a solid-phase assay, we evaluated the binding of these HAs to synthetic glycopolymers. Both TX37-H5N1 and VN1203-H5N1 HAs showed a clear preference for α2-3-linked sialosides (avian-type receptor) over α2-6-linked sialosides (human-type receptor) (Figure 1). TX37-H5N1 HA exhibited preferential binding to sialyl Lewis X (sLeX), a fucosylated α2-3 sialosides, compared to VN1203-H5N1 HA, consistent with findings for other clade 2.3.4.4b H5Nx viruses^11^. These results confirm that both strains retain the typical receptor-binding specificity of avian influenza viruses, with TX37-H5N1 showing some differences in specificity for modified sialic acid structures.

**Figure 1.**
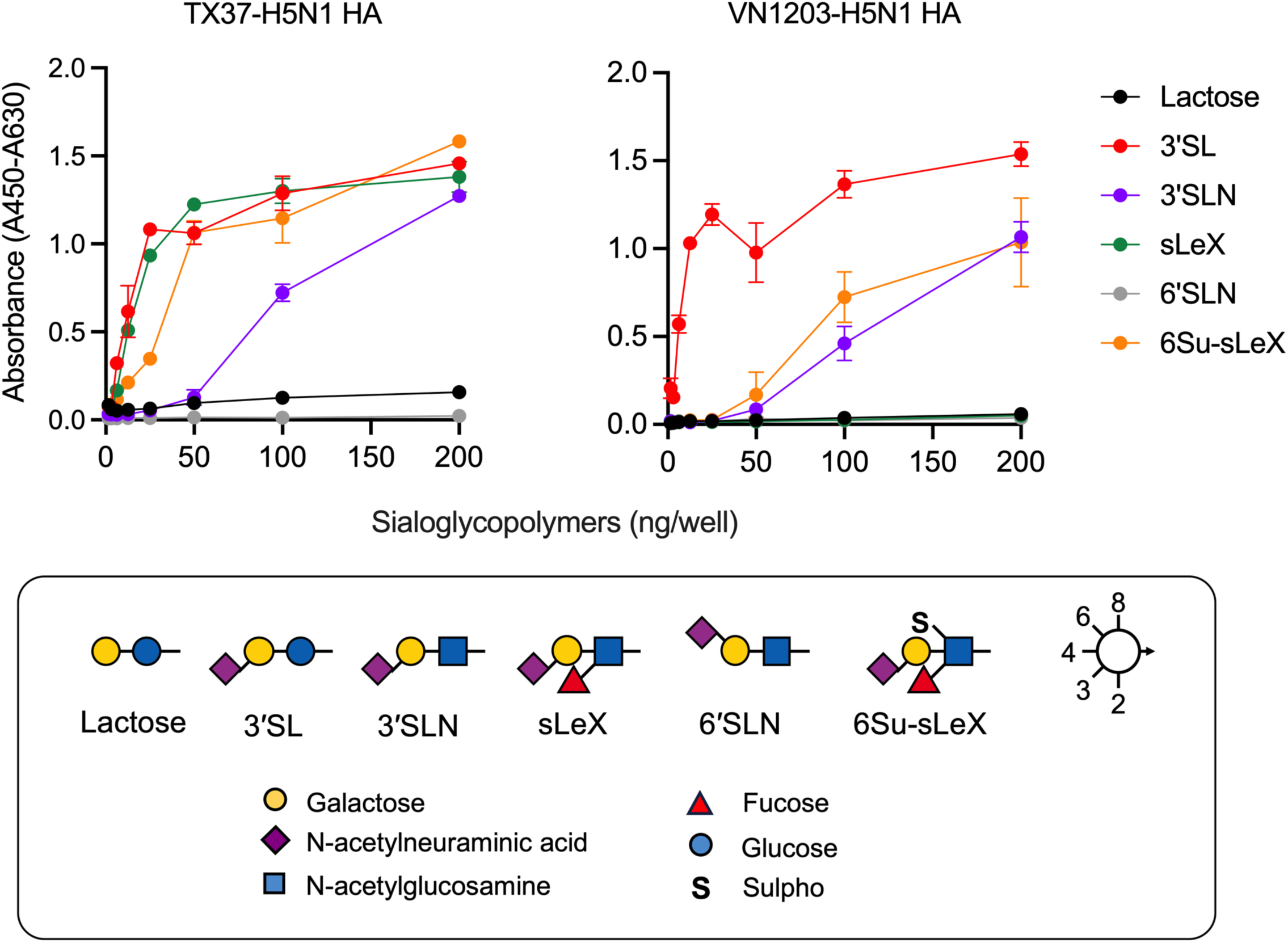
Receptor-binding profiles of influenza virus H5 hemagglutinin. TX37-H5N1 retains avian-type receptor-binding preference, like the conventional H5N1 virus VN1203-H5N1. Binding of recombinant hemagglutinins (HAs) derived from TX37-H5N1 and VN1203-H5N1 to glycopolymers based on poly[N-(2-hydroxyethyl)acrylamide] (PAA) backbones: lactose-PAA (Lactose, black), 3′-sialyllactose-PAA (3′SL, red), 3′-sialyllactosamine-PAA (3′SLN, purple), sialyl Lewis X-PAA (sLeX, green), 6′-sialyllactosamine-PAA (6′SLN, grey), and 6-O-GlcNAc-sulpho-sialyl Lewis X-PAA (6Su-sLex, orange), was evaluated using a solid-phase direct binding assay. Data are presented as the mean ± s.d. (n = 3).

### Human respiratory organoids express receptors for both human and avian influenza viruses

To ensure that our human respiratory organoid model provides a physiologically relevant platform for influenza infection studies, we characterized the expression profile of the sialic acid receptors on the organoid surface. Our iPSC-derived human respiratory organoids contain major airway epithelial cell types, including ciliated cells (Figure 2A). Immunofluorescence staining confirmed the expression of acetylated α-tubulin, a marker for ciliated cells, demonstrating that the apical surfaces were oriented outwards (Figure 2B). We then used a panel of previously characterized recombinant hemagglutinins (HAs) to profile sialoside expression. The probe derived from A/Ezo red fox/Hokkaido/1/2022 (Fox/Hok/1), which broadly binds to α2-3-linked sialosides (avian-type), and the probe from A/New Caledonia/20/1999 (NC/20), targeting α2-6-linked sialosides (human-type), both showed positive binding, indicating that the organoids express both α2-3 Sia and α2-6 Sia receptors. In addition, the probe from A/duck/Mongolia/54/2001 (Dk/Mon), which recognizes unmodified α2-3 Sia, and that from A/chicken/Tainan/V156/1999 (Ck/TN), which binds to sulfated α2-3 Sia, also exhibited clear binding. In contrast, the probe from A/chicken/Ibaraki/1/2005 (Ck/Iba), which recognizes fucosylated α2-3 Sia, showed weaker binding, suggesting that fucosylated α2-3 Sia are expressed at low levels in our respiratory organoids. To confirm the sialic acid dependence of these interactions, organoids were treated with sialidase, which markedly reduced the binding signals of all HA probes (Figure 2C). Finally, HA probes derived from TX37-H5N1 and VN1203-H5N1 demonstrated robust binding to the organoid surface, which was markedly diminished by sialidase treatment, confirming sialic acid-dependent binding (Figure 2D). Together, these results demonstrate that the human respiratory organoids used in this study express both avian-type and human-type receptors, and that the HAs of TX37-H5N1 and VN1203-H5N1 specifically bind to sialosides on the organoid surface. These findings validate the human respiratory organoids as a physiologically relevant model that expresses key viral entry receptors relevant to zoonotic H5N1 infections.

**Figure 2.**
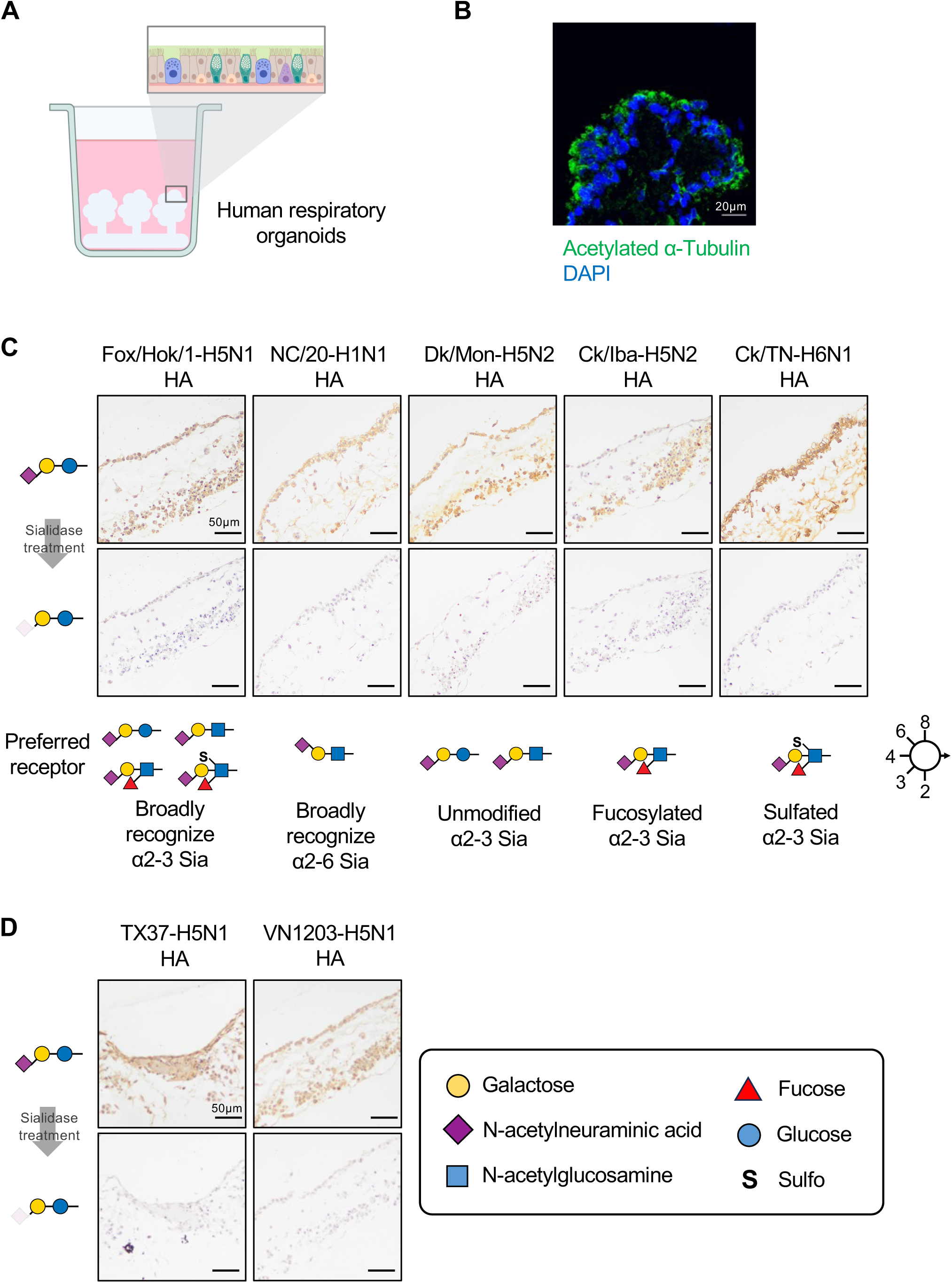
Analysis of sialic acid receptor expression in human respiratory organoids by use of influenza virus HA probes. **(A)** Schematic representation of the experimental design of the human iPSC-derived respiratory organoids. **(B)** Immunofluorescence analysis (IFA) of uninfected respiratory organoids. Acetylated α-tubulin (green), a marker of ciliated cells, was detected in human iPSC-derived respiratory organoids. Nuclei were counterstained with DAPI (blue). Scale bar, 20 μm. **(C)** Recombinant hemagglutinins (HAs) with distinct glycan-binding specificities were used to determine the receptor profile of the human respiratory organoids: Fox/Hok/1-H5N1 (A/Ezo red fox/Hokkaido/1/2022) HA broadly recognizes α2-3-linked sialosides (Sia); NC/20-H1N1 (A/New Caledonia/20/1999) HA broadly recognizes α2-6 Sia; Dk/Mon-H5N2 (A/duck/Mongolia/54/2001) HA preferentially recognizes unmodified α2-3 Sia; Ck/Iba-H5N2 (A/chicken/Ibaraki/1/2005) HA recognizes fucosylated α2-3 Sia; and Ck/TN-H6N1 (A/chicken/Tainan/V156/1999) HA recognizes sulphated α2-3 Sia. Sialidase-treated negative controls are shown in the lower panels. Scale bar, 50 μm. **(D)** Binding of recombinant HAs from TX37-H5N1 and VN1203-H5N1 to human respiratory organoids. Sialidase-treated negative controls are shown in the lower panels. Scale bar, 50 μm.

### TX37-H5N1 and VN1203-H5N1 replicate efficiently in human respiratory organoids

Given the discrepancy between animal model pathogenicity and the mild symptoms experienced by humans upon infection with B3.13 viruses, we hypothesized that virus replication efficiency might differ in human respiratory tissues compared with those of animal models. Therefore, we infected the human respiratory organoids with TX37-H5N1, VN1203-H5N1, and the pandemic human H1N1 influenza virus A/California/04/2009 (Cal04-H1N1) at a dose of 8 × 10^4^ plaque-forming unit (PFU). Viral titers in culture supernatants were measured over time (Figure 3A). TX37-H5N1 grew rapidly and achieved titers of 9.12 ± 0.04 log10 PFU/mL at 48 h post-infection (h.p.i.). VN1203-H5N1 exhibited similarly rapid replication, reaching 8.55 ± 0.03 log10 PFU/mL at 48 h.p.i. Notably, TX37-H5N1 reached higher titers than VN1203-H5N1 at 24, 48, and 72 h.p.i. (*p*<0.01 at each time point), suggesting more efficient replication throughout the infection course. In contrast, Cal04-H1N1 showed a slower replication profile, with viral loads gradually increasing and peaking at 5.95 ± 0.29 log10 PFU/mL at 72 h.p.i. Similar dose-dependent trends were observed across the other inocula tested, with TX37-H5N1 and VN1203-H5N1 showing comparable replication at lower doses (8 × 10^3^ and 8 × 10^2^ PFU), and both H5N1 strains consistently replicating more efficiently than Cal04-H1N1 across all doses tested (Figure 3A). Immunohistochemical staining of organoid sections at 72 h.p.i. revealed striking differences in infection patterns: TX37-H5N1 and VN1203-H5N1 infections resulted in widespread and intense influenza NP antigen staining, whereas Cal04-H1N1 infection was restricted to scattered positive cells (Figure 3B). These results indicate that both H5N1 strains replicate more efficiently than Cal04-H1N1 in this organoid model, with TX37-H5N1 exhibiting the greatest replicative ability.

**Figure 3.**
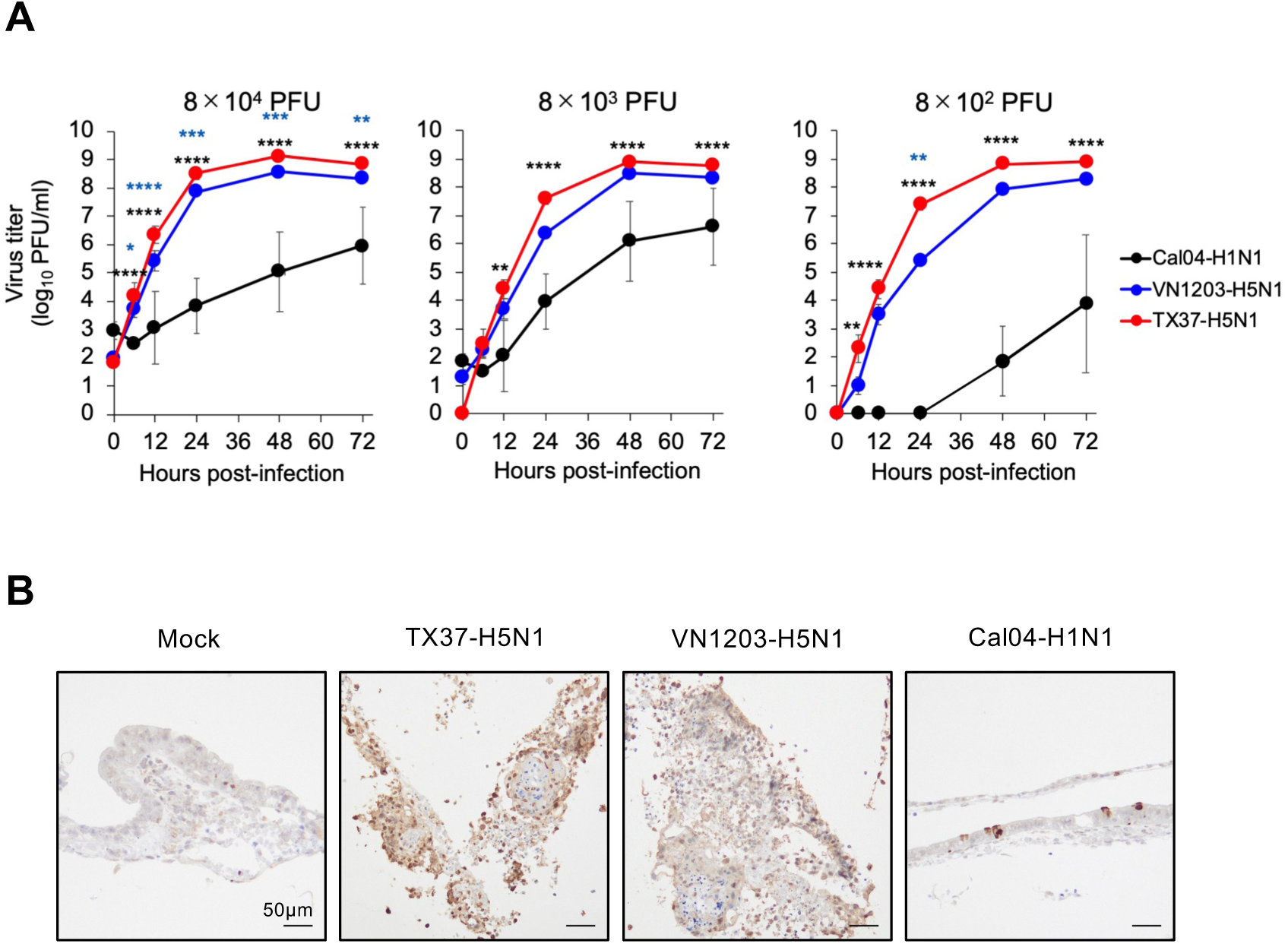
Viral replication kinetics and detection of viral antigen in human respiratory organoids. **(A)** Growth kinetics of TX37-H5N1, VN1203-H5N1, and Cal04-H1N1 viruses at a dose of 8 × 10^4^, 8 × 10³, or 8 × 10^2^ PFU in human respiratory organoids. Culture supernatants were collected at 0, 6, 12, 24, 48, and 72 h.p.i., and virus titers were determined by plaque assay in MDCK cells. Data are presented as the mean ± s.d. (n = 3). Statistical analysis was performed using a two-way ANOVA followed by Dunnett’s multiple comparisons test. At each timepoint, black asterisk denote comparisons of TX37-H5N1 to Cal04-H1N1 and blue asterisk denote comparisons of TX37-H5N1 to VN1203-H5N1. \**p*<0.05, \*\**p*<0.01, \*\*\**p*<0.001, \*\*\*\**p*<0.0001. **(B)** Immunohistochemical detection of NP antigens in infected respiratory organoids. Scale bar, 50 μm.

### Assessment of anti-influenza drug efficacy in human respiratory organoids

To demonstrate the utility of our human respiratory organoid model for antiviral drug assessment, we evaluated the efficacy of three FDA-approved influenza antivirals with distinct mechanisms of action: oseltamivir (a neuraminidase inhibitor), favipiravir (a viral RNA polymerase inhibitor), and baloxavir (a cap-dependent endonuclease inhibitor). Organoids were infected with standardized doses (200 PFU for H5N1 strains; 100,000 PFU for Cal04-H1N1) and treated with clinically relevant drug concentrations (10 μM oseltamivir, 20 μg/mL favipiravir, 40 nM baloxavir). Viral titers were measured at 72 h.p.i.

In Cal04-H1N1-infected organoids, oseltamivir and baloxavir suppressed viral replication to undetectable levels, whereas favipiravir had little effect. For both H5N1 strains, oseltamivir achieved significant virus suppression, reducing TX37-H5N1 titers by 3.17 log (from 8.99 to 5.82 log10 PFU/mL, *p*=0.0004) and VN1203-H5N1 titers by 3.07 log (from 7.75 to 4.68 log10 PFU/mL, *p*<0.0001), representing approximately 1,000-fold reductions (Figure 4). Baloxavir demonstrated superior efficacy against both H5N1 strains, achieving near-complete suppression with >6.75-log reductions for VN1203-H5N1 and >7.35-log reductions for TX37-H5N1 compared to the control group. In contrast, favipiravir showed strain-specific effects: while largely ineffective against VN1203-H5N1 and Cal04-H1N1, it achieved modest but statistically significant inhibitory activity against TX37-H5N1 (0.77 log reduction, *p*=0.0014), suggesting that baloxavir has strong antiviral activity against these three viral strains, oseltamivir is effective against these strains but less effective against TX37-H5N1 and VN1203-H5N1 than against Cal04-H1N1, and favipiravir has limited efficacy in this organoid model.

**Figure 4.**
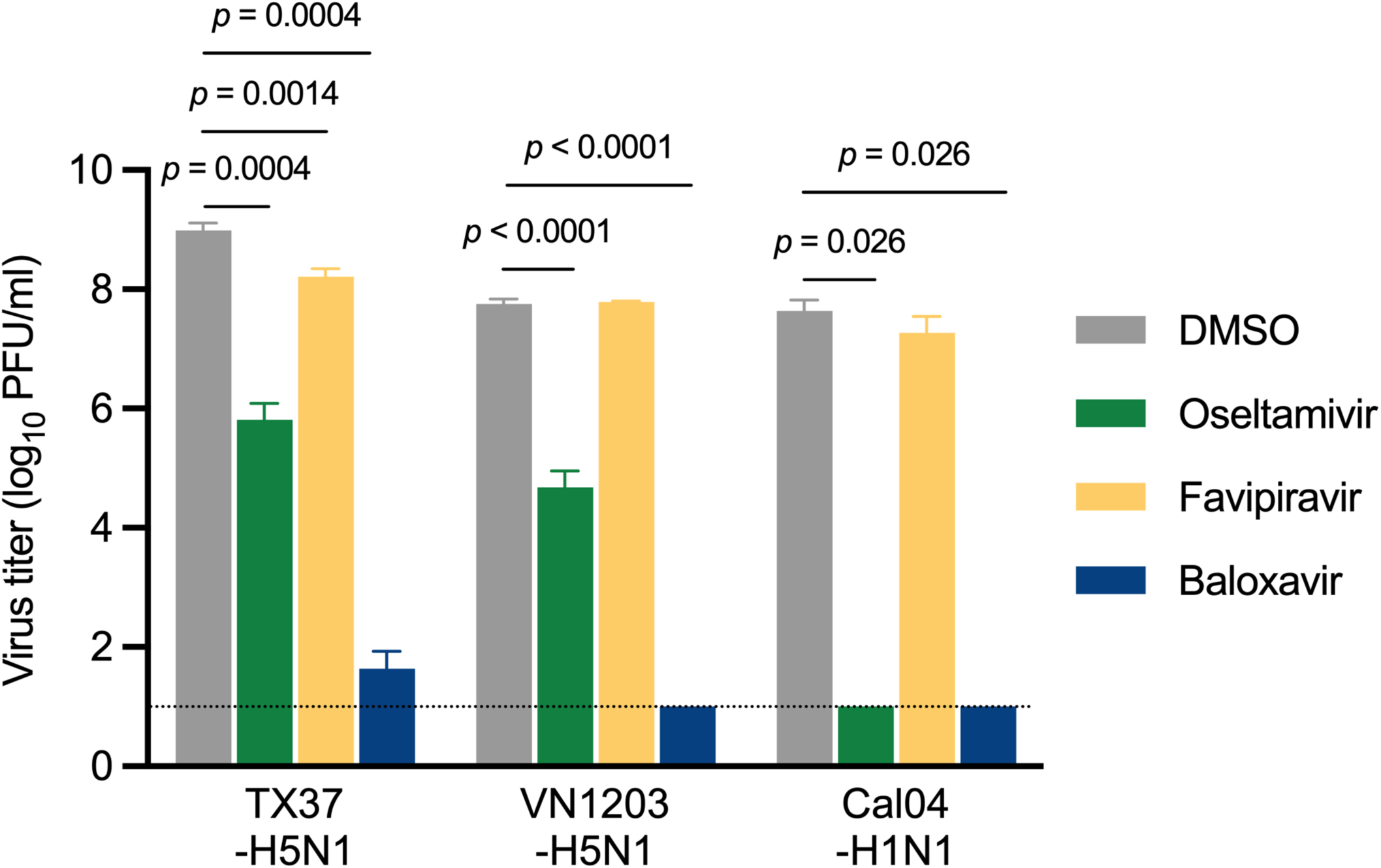
Efficacy of influenza antiviral drugs in human respiratory organoids. Antiviral drug sensitivity of viruses in respiratory organoids. Respiratory organoids were infected with 200 PFU of VN1203-H5N1 or TX37-H5N1 or 100,000 PFU of Cal04-H1N1. Infected cells were incubated in the presence of DMSO, 10 µM oseltamivir acid, 20 µg/mL favipiravir, or 40 nM baloxavir acid. Culture supernatants were collected at 72 h.p.i., and virus titers were determined by plaque assay on MDCK cells. Data are presented as the mean ± s.d. (n = 3). Statistical analysis was performed using a one-way ANOVA followed by Tukey’s multiple comparisons test. The dotted line indicates the detection limit (1.0 log10 PFU/ml).

These results confirm that our human respiratory organoid system can be used to assess antiviral drug efficacies with sensitivity comparable to traditional cell culture systems while providing a more physiologically relevant platform for evaluating therapeutic responses under near-physiological conditions.

### TX37-H5N1 induces minimal cytokine responses despite efficient replication in human respiratory organoids

Given the striking difference in clinical outcomes between TX37-H5N1 (mild symptoms) and historical H5N1 strains (severe disease), we hypothesized that innate immune responses might differ between these viruses. To test this hypothesis, we measured cytokine and chemokine levels in the culture supernatants of virus-infected respiratory organoids at 72 h.p.i., focusing on key mediators of antiviral and inflammatory responses (Figure 5). We used the LEGENDplex Human Anti-Virus Response Panel, which allows simultaneous quantification of 13 human proteins, including type I interferons (IFN-α2, IFN-β), type II interferons (IFN-γ), type III interferons (IFN-λ1, IFN-λ2), pro-inflammatory cytokines (IL-1β, IL-6, TNF-α), chemokines (IL-8, IP-10), and immunoregulatory factors (IL-10, IL-12p70, GM-CSF). VN1203-H5N1 infection induced robust production of multiple cytokines, with IFN-β levels reaching 280.2 ± 43.1 pg/mL, IL-6 concentrations of 171.4 ± 46.2 pg/mL, IL-8 levels of 920.3 ± 211.9 pg/mL, and IP-10 concentrations of 264.7 ± 40.5 pg/mL (Figure 5). This cytokine profile indicates a robust inflammatory response typically associated with H5N1 infection and consistent with the elevated cytokine levels reported in severe cases^10^. In contrast, both TX37-H5N1 and Cal04-H1N1 infections elicited markedly attenuated responses across all measured cytokines, and levels of IFN-β, IL-6, and IP-10 were significantly lower compared to those induced by VN1203-H5N1 (Figure 5). Although IL-8 was detectable in the TX37-H5N1- and Cal04-H1N1-infected organoids, its level was significantly lower than that in the VN1203-H5N1-infected organoids.

**Figure 5.**
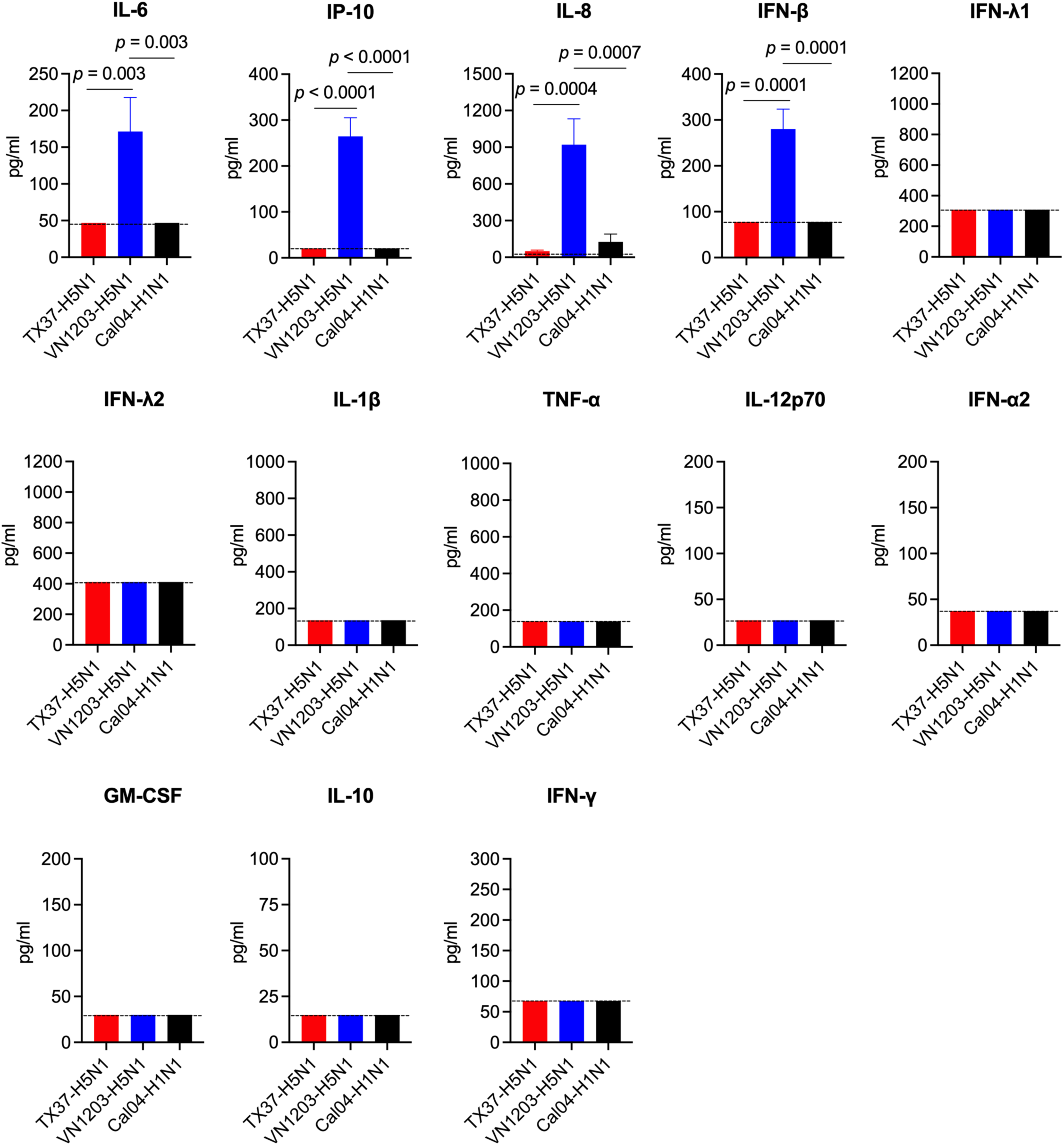
Cytokine and interferon responses in virus-infected respiratory organoids. Quantification of IFN-λ1, IL-1β, IL-6, TNF-α, IP-10, IL-8, IL-12p70, IFN-α2, IFN-λ2, GM-CSF, IFN-β, IL-10, and IFN-γ protein levels in the supernatants of virus-infected respiratory organoids at 72 h.p.i. Data are presented as the mean ± s.d. (n = 3). Statistical analysis was performed using a one-way ANOVA followed by Tukey’s multiple comparisons test. In all panels, the dotted line denotes the assay detection limit.

These findings demonstrate that, despite achieving higher viral titers than VN1203-H5N1, TX37-H5N1 triggers substantially weaker interferon and inflammatory cytokine responses. This pattern suggests an active immune evasion mechanism that may contribute to the mild clinical presentation observed in human B3.13 infections, contrasting sharply with the pathogenic hyperinflammatory responses associated with historic H5N1 strains.

### TX37-H5N1 induces selective suppression of antiviral programs

To investigate the temporal dynamics of host antiviral responses, we performed bulk RNA-seq on human respiratory organoids infected with TX37-H5N1, VN1203-H5N1, or Cal04-H1N1 at 12, 24, and 48 h.p.i. (Figure 6A). Principal component analysis (PCA) showed that transcriptomes were separated by both time and viral strain (Figure 6B), with PC1 accounting for 62% of the variance and correlating with time post-infection, whereas PC2 separated samples primarily by viral strain. Differential gene expression analysis revealed that at 12 h.p.i., only a few mRNAs were significantly altered compared to the mock-infected control, with 2, 6, and 4 differentially expressed genes (DEGs) detected in TX37-H5N1, VN1203-H5N1, and Cal04-H1N1 infections, respectively (Figure 6C). By 48 h.p.i., however, TX37-H5N1 and VN1203-H5N1 infections induced extensive transcriptional changes, with more than 5,000 DEGs identified in each group, whereas Cal04-H1N1 infection continued to induce limited changes. More than 50% of the DEGs were shared between TX37-H5N1 and VN1203-H5N1 (Figure 6C), reflecting a common H5N1-induced host response. Each strain also triggered unique gene expression patterns (Supplementary Table 1). Given the minimal changes in the Cal04-H1N1 group, our subsequent analyses focused on the H5N1 viruses.

**Figure 6.**
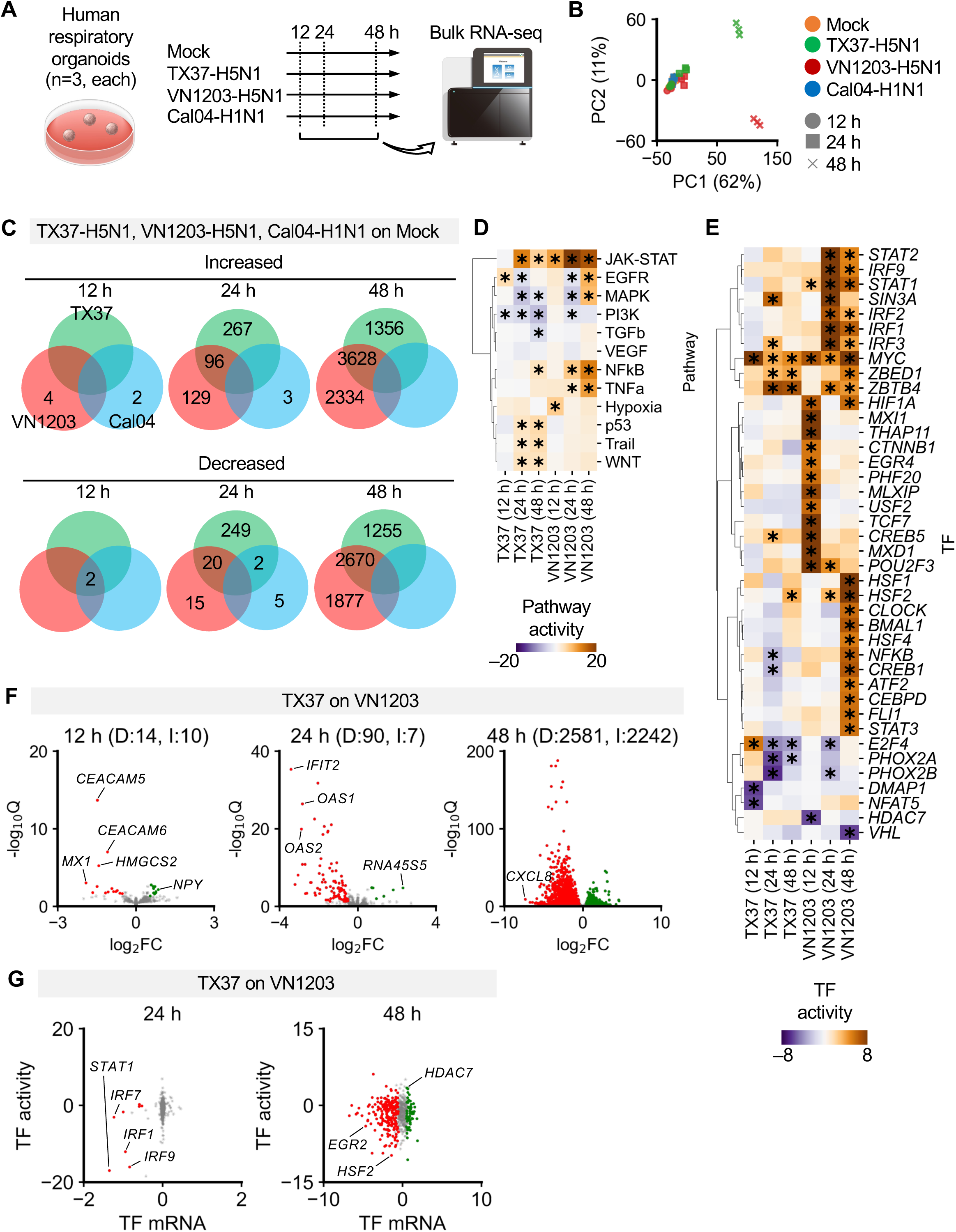
Antiviral response profiling by bulk RNA-seq. **(A)** Schematic overview of the experimental workflow. Human respiratory organoids were infected with influenza virus strains TX37-H5N1, VN1203-H5N1, Cal04-H1N1, or Mock-treated. Total RNA was collected at 12, 24, and 48 h.p.i. and subjected to bulk RNA-seq. **(B)** Principal component analysis (PCA) of the transcriptomic profiles for each condition. Each point represents an individual sample (n = 3 per condition per time point). **(C)** Venn diagrams showing the number of differentially expressed genes (DEGs; defined as FDR < 0.05, | log2(Fold Change) | > 0.5 compared to mock-treated controls) identified at each time point for each viral strain. Only non-zero values are shown. **(D)** Heatmap showing pathway activity inferred using PROGENy and decoupleR. Rows correspond to individual pathways, while columns indicate each condition and time point. Color scale indicates pathway activity values. *Adjusted *p*-values < 0.01. **(E)** Heatmap of transcription factor activity inferred by CollecTRI and decoupleR. Rows correspond to individual transcription factors (TFs), while columns indicate each condition and time point. Color scale indicates TF activity values. TFs with at least one condition exhibiting an absolute activity value ≥ 5 are shown. *Adjusted *p*-values < 0.01. **(F)** Volcano plots comparing transcriptomic changes between TX37-H5N1 and VN1203-H5N1 infections at each time point. mRNAs with FDR < 0.05 and | log2(Fold Change) | > 0.5 were considered significant; “I” and “D” indicate the number of mRNAs significantly increased or decreased in TX37-H5N1 compared to VN1203-H5N1, respectively. **(G)** Scatter plots comparing TF activity inferred by CollecTRI and decoupleR (y-axis) and corresponding relative TF mRNA abundance of TX37-H5N1 compared to VN1203-H5N1 [x-axis, log2(Fold Change)] at 24 and 48 h.p.i. Each dot represents a single TF.

Pathway and transcription factor (TF) activity analysis revealed sequential activation of antiviral and inflammatory signaling cascades in the VN1203-H5N1-infected organoids. JAK-STAT and hypoxia pathways were activated as early as 12 h.p.i., followed by pronounced NFκB and TNFα signaling at 48 h.p.i. (Figure 6D). This was accompanied by sequential upregulation of TFs, including HIF1A, STAT1, STAT2, and IRFs, and subsequent activation of STAT3, NFκB, and heat shock factors (Figure 6E). In contrast, these signaling and TF responses were largely absent or markedly attenuated in TX37-H5N1-infected organoids throughout the time course. The limited activation of these pathways and TFs in TX37-H5N1-infected organoids was reflected in weak induction of downstream target genes, including *IFNB*, *CXCL8,* and *CXCL10,* which are canonical interferon-stimulated genes (ISGs) regulated by STAT1 and STAT2. At the protein level, cytokine measurements similarly revealed reduced production of IFN-β, IL-8 and IP-10 in TX37-H5N1-infected organoids compared to VN1203-H5N1-infected organoids (Figure 5).

Direct comparison between TX37-H5N1 and VN1203-H5N1 confirmed that VN1203-H5N1 more potently upregulated antiviral and ISG-related transcripts, including *MX1*, *IFIT2*, and *OAS1/2*, across time points (Figure 6F). Comparative analysis of TF activity revealed a greater induction of both mRNA expression and TF activity of *STAT1* and *IRF1* in VN1203-H5N1-infected organoids at 24 h.p.i. compared to in TX37-H5N1-infected organoids (Figure 6G). Collectively, these results indicate that while both TX37-H5N1 and VN1203-H5N1 induce transcriptomic changes, the antiviral and inflammatory responses are significantly attenuated in TX37-H5N1-infected organoids, suggesting that TX37-H5N1 infection fails to initiate effective activation of canonical antiviral transcriptional programs, particularly the STAT-IRF signaling axis, despite robust viral replication.

### Single-cell analysis confirms cell type-independent STAT-IRF suppression by TX37-H5N1

To investigate cell-type-specific host responses and viral distribution, we performed fixed single-cell RNA profiling of human respiratory organoids at 24 h.p.i. with TX37-H5N1, VN1203-H5N1, or Cal04-H1N1 (Figure 7A). Uniform Manifold Approximation and Projection (UMAP) visualization of the transcriptomic data indicated that the Mock and Cal04-H1N1 samples had largely overlapping distributions, whereas organoids infected with TX37-H5N1 or VN1203-H5N1 induced significant shifts in global gene expression (Figure 7B, upper panel), consistent with the results of bulk RNA-seq (Figure 6). Cell-type annotation showed a relative expansion of alveolar fibroblasts and depletion of epithelial progenitor cells, particularly mid-stalk cells (Figure 7B, lower panel; Figure 7C). Mid-stalk cells represent transient epithelial progenitors that arise during fetal lung development and differentiate into alveolar epithelial type 1 (AT1) and AT2 cells^12^. This shift in progenitor composition was accompanied by epithelial damage, as observed by IHC (Figure 3B). TX37-H5N1-infected organoids exhibited a high proportion of infected cells and widespread viral distribution across multiple cell types, in contrast to the more limited infection observed in the VN1203-H5N1 and Cal04-H1N1 samples (Figure 7D, E).

**Figure 7.**
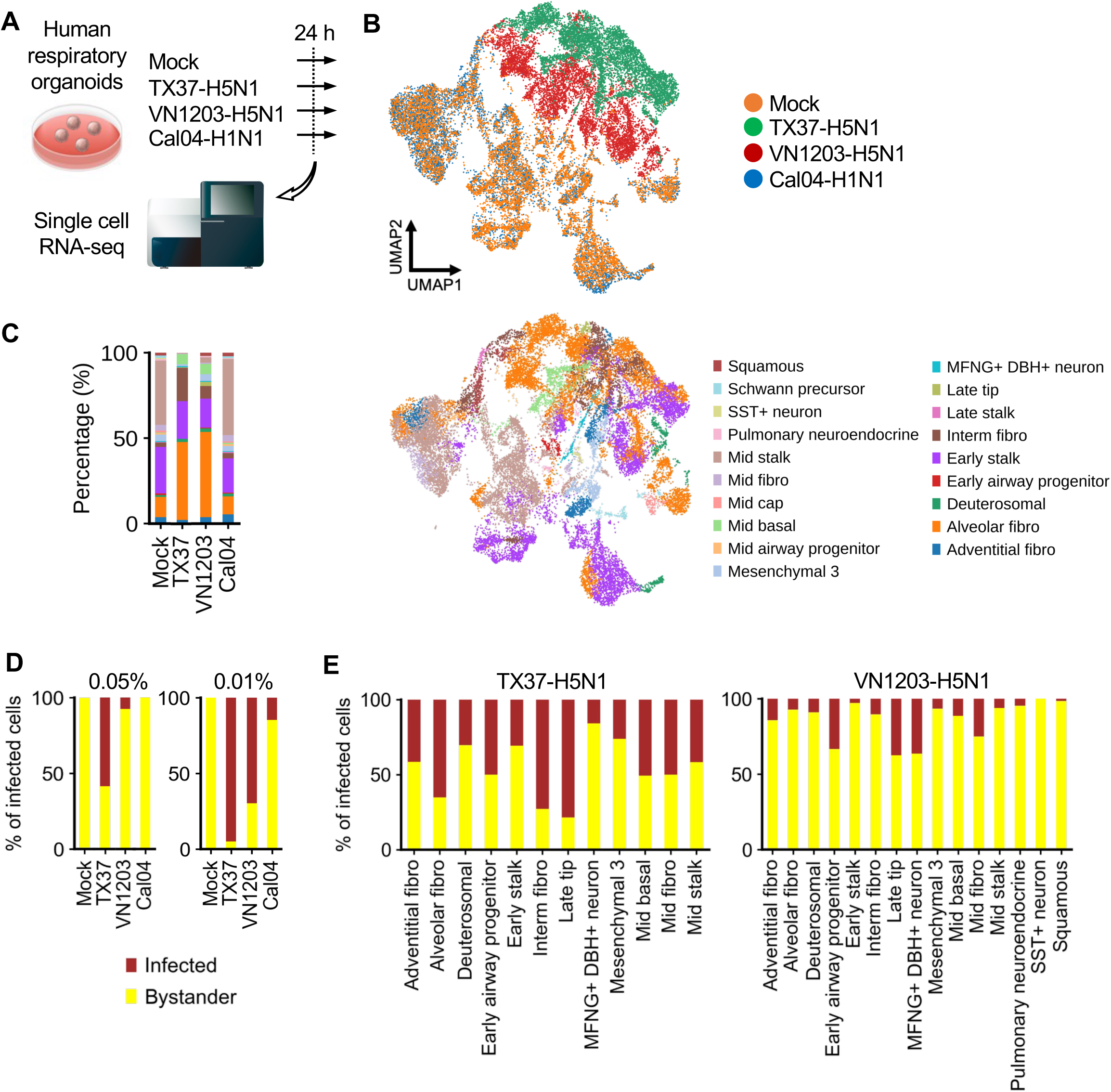
Single-cell transcriptomic landscape and infection prevalence in respiratory organoids. **(A)** Schematic representation of the experimental procedure. Human respiratory organoids were infected with the influenza virus strains TX37-H5N1, VN1203-H5N1, Cal04-H1N1, or Mock-treated. At 24 h.p.i., organoids were fixed with formaldehyde and processed into single-cell suspensions according to the Fixation of Cells & Nuclei for Chromium Fixed RNA Profiling protocol for subsequent single-cell RNA sequencing. **(B)** UMAP visualization of single cells, colored according to infection condition (Mock, TX37-H5N1, VN1203-H5N1, Cal04-H1N1; Upper) or by annotated cell type (lower). **(C)** Bar plot showing the proportion of each cell type within each condition. Colors correspond to cell types in Figure 7B. **(D)** Prevalence of infected cells under each condition. Cells were classified as infected when viral transcript counts exceeded 0.05% or 0.01% of the total host transcript counts per cell. **(E)** Prevalence of infected cells with viral transcript counts exceeded 0.05% in each annotated cell type for TX37-H5N1 and VN1203-H5N1 infections.

In TX37-H5N1-infected cells, viral transcript abundance was positively correlated with stress-response genes such as *HSPA1B*, *BAG3*, and *SNHG32*, and negatively correlated with genes involved in cell adhesion and extracellular matrix formation (*COL1A2* and *WLS*) (Figure 8A). In contrast, VN1203-H5N1-infected cells showed positive correlations between viral load and genes associated with protein synthesis and cytoskeletal remodeling (*EEF1G*, *CFL1*, *TMSB10*, *TMSB4X*), whereas transcriptional and developmental regulators (*SOX2*, *MSX1, ZIC1*) were suppressed. TF activity analysis revealed that while VN1203-H5N1 infection induced strong activation of canonical antiviral regulators, including *STAT1*, *STAT2*, *IRF1*, and *IRF2* (Figure 8B), these responses were mostly absent in TX37-H5N1-infected cells. Instead, TX37-H5N1 especially upregulated *HSF2* and *SMAD9* (Figure 8B). The lack of STAT-IRF activity in TX37-H5N1-infected cells corresponded to low transcript levels of these factors (Figure 8C, D), consistent with the bulk RNA-seq data (Figure 6). Together, these results suggest that TX37-H5N1 infection, despite eliciting high viral loads and widespread gene expression changes, leads to limited activation of antiviral responses, particularly those involving STAT and IRF family members.

**Figure 8.**
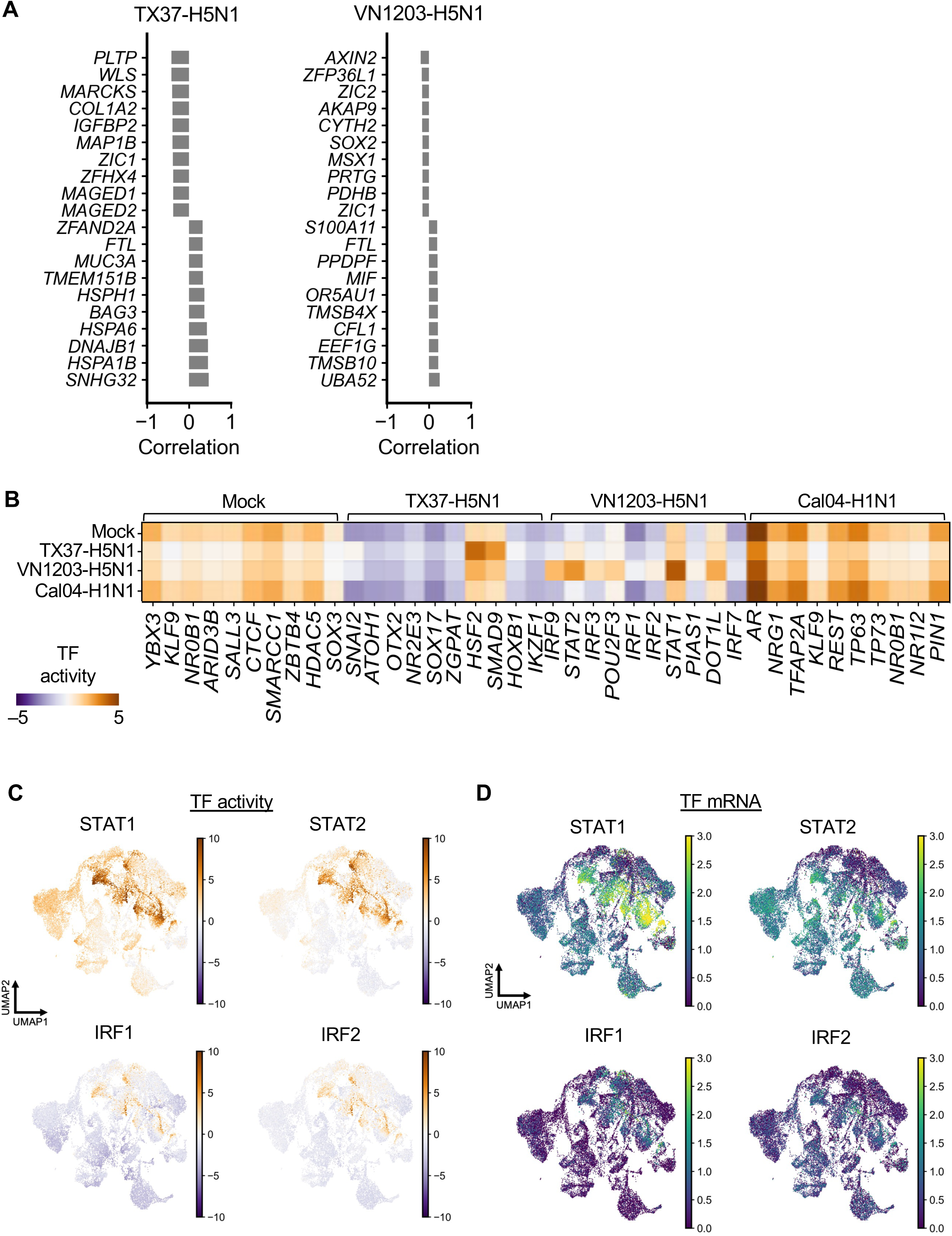
Gene expression correlations and transcription factor activity in virus-infected cells. **(A)** Correlation coefficients between viral transcript abundance and host mRNA expression in TX37-H5N1- and VN1203-H5N1-infected cells. The top 10 genes with the highest positive and negative correlation coefficients are shown for each infection. All mRNAs shown have adjusted *p*-values < 0.05. **(B)** Heatmap showing inferred TF activity calculated by CollecTRI and decoupleR across conditions. The 10 transcription factors with the highest differential activity compared to other conditions are shown for each condition. **(C)** UMAP visualization of single cells colored by inferred TF activity for *STAT1*, *STAT2*, *IRF1,* and *IRF2*. **(D)** UMAP visualization of single cells colored by mRNA expression levels of *STAT1*, *STAT2*, *IRF1,* and *IRF2*.

## Discussion

The H5N1 HPAIV has caused sporadic but severe infections in humans. Early human cases in the 2000s were characterized by viral pneumonia, ARDS, and excessive cytokine production, with a reported case fatality rate of approximately 48%^5^. Viral strains from that era were also highly lethal in animal models such as mice and ferrets, demonstrating high pathogenicity^7,8^. In contrast, the B3.13 genotype of clade 2.3.4.4b H5N1 virus (TX37-H5N1), first confirmed in a human case in the United States in 2024^3^, retains high pathogenicity in animal models but causes only mild symptoms such as conjunctivitis or upper respiratory illness in humans^4,7,8^. The basis for this striking difference between experimental models and human clinical presentation remains unknown. In this study, we used human iPSC-derived respiratory organoids to compare a previously characterized H5N1 human isolate (VN1203-H5N1) and the recently identified cattle-derived human isolate TX37-H5N1. The replication efficiency of TX37-H5N1 was comparable to or greater than that of VN1203-H5N1 (Figure 3), but TX37-H5N1 better suppressed innate immune responses (Figure 5, 6), particularly with respect to the activation of the STAT–IRF axis (Figure 7, 8). These findings suggest that suppression of innate immunity may contribute to the attenuated pathogenicity of B3.13 viruses in humans and highlight the importance of innate immune modulation in determining H5N1 virulence.

The human respiratory organoid model employed in this study mirrors physiologic virus-host interactions through its inclusion of airway epithelial cells, fibroblasts, and immune-related lineages in a three-dimensional structure. This model has previously been used to analyze host responses to SARS-CoV-2^13,14^ and respiratory syncytial virus^15^, demonstrating its capacity to capture human-specific infection dynamics and immune responses. In the present study, the organoid model enabled us to uncover the molecular basis for the mild clinical presentation of TX37-H5N1, despite its high lethality in animal models. In previous animal model studies, H5N1 virus infection often triggered hyperinflammatory responses characterized by elevated levels of pro-inflammatory cytokines^16^. Although cytokine levels were not directly measured in TX37-infected animals, the high lethality observed in these models suggests a similar immunopathological response may have occurred. In contrast, our human respiratory organoid system revealed a marked suppression of innate immune signaling following TX37-H5N1 infection (Figure 6 – 8). These results underscore the strength of this organoid system for visualizing species-specific host responses and evaluating subtle differences in innate immune regulation that may be missed in traditional animal models. Our findings support the use of human organoids as a complementary platform for analyzing discrepancies between experimental and clinical observations. Furthermore, we demonstrated the applicability of this model for evaluating antiviral efficacy. TX37-H5N1 was sensitive to baloxavir and oseltamivir but showed limited response to favipiravir, reinforcing the feasibility of using this system to assess viral replication and drug response under near-physiological conditions.

As efforts to restrict animal experimentation gain momentum, driven by the principles of the 3Rs (Replacement, Reduction, Refinement), human-derived organoid platforms could play a more central role in infectious disease research and therapeutic evaluation^17^. Nevertheless, limitations remain. The model we used is well-suited for analyzing innate responses but cannot mimic adaptive immunity (e.g., T and B cell interactions), systemic immune networks, or the contributions of vascular and neural components. Moreover, scRNA-seq analysis indicates that organoids primarily comprise immature, fetal-like cell populations, which may limit their ability to fully model adult tissue responses. The model also does not capture post-infection regeneration or the full pharmacokinetics of antiviral drugs. Addressing these limitations requires integration with clinical datasets and complementary use of *in vivo* models. Advances such as co-culture with immune and endothelial cells or organ-on-a-chip technologies^18,19^ may help enhance the physiological relevance of the model.

In response to viral infection, innate immune sensors such as pattern recognition receptors (PRRs) activate key transcription factors including the STAT (Signal Transducer and Activator of Transcription) and IRF (Interferon Regulatory Factor) families, leading to the induction of type I interferons and inflammatory cytokines^20^. These responses establish an antiviral state and recruit and activate immune cells to fight the infection^21^. However, excessive activation of the STAT–IRF axis can lead to immunopathology^22^. Indeed, VN1203-H5N1 infection strongly upregulated this axis, leading to rapid expression of interferon-stimulated and proinflammatory genes, which are hallmarks of the cytokine storm and ARDS observed in severe H5N1 cases^10^. In contrast, despite robust replication, TX37-H5N1 failed to activate the STAT–IRF axis (Figure 6). Notably, the transcription factor activities of key mediators such as STAT1, STAT2, IRF1, and IRF2 were not markedly stimulated in the TX37-H5N1 respiratory organoids, and expression of antiviral effector genes was markedly suppressed even in infected cells (Figure 7, 8). Consistent with these findings, cytokine quantification demonstrated attenuated secretion of IFN-β and IP-10, as well as reduced levels of IL-6 and IL-8, supporting the notion that suppression of the STAT–IRF axis diminishes both antiviral and inflammatory responses (Figure 5). This “stealth infection” phenotype suggests that immune evasion via innate suppression may underlie the low pathogenicity of TX37-H5N1. Our findings further support the idea that the extent of STAT–IRF activation influences H5N1 virulence. Elucidating the regulatory mechanisms of this pathway and identifying viral factors (e.g., NS1) responsible for its suppression may yield new insights into pathogenesis and inform the development of host-targeted therapies.

Our study provides evidence that selective suppression of innate immunity may explain the mild clinical manifestations of B3.13 H5N1 viruses in humans and offers new perspectives for reconciling inconsistencies between animal models and clinical outcomes. Viruses like TX37-H5N1 that replicate efficiently yet cause only mild disease in humans may evade detection if only traditional risk assessment strategies are employed (e.g., measuring LD₅₀ values and viral titers in mice and ferrets). Immune response profiling using human-derived organoids, as demonstrated here, offers a powerful complementary approach to capture human-specific innate immune responses that may not align with animal-model outcomes. Moreover, viruses with low-pathogenic but high-replication phenotypes pose unique challenges for public health. Such viruses may silently spread through asymptomatic infection and prolonged viral shedding. While no sustained human-to-human transmission of B3.13 viruses has been reported to date^23^, future mutations may alter transmissibility. Because no single model can fully capture public health threats of emerging influenza viruses, an integrated framework is needed to evaluate viral replication efficiency, immune modulation, and transmissibility. Finally, the suppression of the STAT–IRF pathway identified in this study may offer new avenues for therapeutic intervention and have implications for pandemic preparedness and emerging infectious disease control. Future efforts should focus on characterizing host immune responses in human cases and performing cross-strain comparisons to identify predictive markers of human pathogenicity.

## Materials and Methods

### Recombinant hemagglutinin production

Complementary DNAs encoding the hemagglutinin (HA) genes of A/Texas/37/2024 (TX37-H5N1), A/Vietnam/1203/2004 (VN1203-H5N1), A/Ezo red fox/Hokkaido/1/2022 (Fox/Hok/1-H5N1), A/New Caledonia/20/1999 (NC/20-H1N1), A/duck/Mongolia/54/2001 (Dk/Mon-H5N2), A/chicken/Ibaraki/1/2005 (Ck/Iba-H5N2), and A/chicken/Tainan/V156/1999 (Ck/TN-H6N1) were cloned into pCD5 expression vectors, as described previously^24,25,26^. In these vectors, the HA genes, lacking transmembrane domains, were fused to a GCN4IL trimerization motif and a Strep-tag II (WSHPQFEK; IBA Lifesciences GmbH, Göttingen, Germany). Recombinant HA (rHA) proteins were expressed in HEK 293S GnTI–/– cells and purified from culture supernatants as previously described^27,28^. Briefly, rHA proteins were captured using Strep-Tactin Sepharose (50% suspension; IBA Lifesciences) at 4°C for 15 min and eluted using an elution buffer containing 1 mM EDTA and 5 mM desthiobiotin (IBA Lifesciences) in phosphate-buffered saline (PBS).

### Solid-phase receptor-binding assay

The receptor-binding specificity of the rHA was assessed using a solid-phase direct binding assay with glycopolymers based on poly[N-(2-hydroxyethyl)acrylamide] (PAA) backbones: lactose-PAA (Lactose), 3′-sialyllactose-PAA (3′SL), 3′-sialyllactosamine-PAA (3′SLN), sialyl Lewis X-PAA (sLeX), 6′-sialyllactosamine-PAA (6′SLN), and 6-O-GlcNAc-sulpho-sialyl Lewis X-PAA (6Su-sLex) (GlycoNZ, Auckland, New Zealand). Each sialylated glycan was serially diluted from 1.56 µM to 200 µM and applied to wells of a Nunc Immobilizer Amino F8 96-well strip well microplate (Thermo Fisher Scientific Inc, Waltham, MA, USA). The plates were then incubated at 37°C for 1 h. Each well was then blocked with PBS with Tween 20 (PBST) containing 2% bovine serum albumin (BSA) at 23–28°C for 1 h. The rHAs were used at 5.0 μg/mL, pre-complexed with horseradish peroxidase (HRP)-linked anti-Strep-tag II mouse antibody (IBA Lifesciences; 1:2000 dilution) and goat anti-mouse IgG-HRP conjugate (Bio-Rad Laboratories, Inc., Hercules, CA, USA; 1:1000 dilution) before incubation for 30 min on ice in PBST containing 0.5% BSA. After the plates were washed, complexes were added and incubated at 23–28°C for 1 h. After an additional wash, 100 μL of 3,3′- tetramethylbenzidine and 0.04% H2O2 was added to each well. The reactions were stopped by adding 50 μL of 2N H2SO4, and absorbance at 450/630 nm was measured using a MULTISKAN JX (Thermo Fisher Scientific). Data represent the mean ± s.d. (n = 3) from a single experiment.

### Lectin histochemistry using recombinant HAs

A panel of previously characterized trimetric recombinant HA (rHA) probes was used to profile sialoside expression in the human respiratory organoids^28^. The rHAs included the following: a broadly α2-3 sialic acid (Sia)-binding probe derived from Fox/Hok/1-H5N1; an α2-6 Sia-specific probe from NC/20-H1N1; a probe targeting unmodified α2-3 Sia from Dk/Mon-H5N2; a fucosylated α2-3 Sia-binding probe from Ck/Iba-H5N2; and a sulphated α2-3 Sia-binding probe from Ck/TN-H6N1. In addition, rHA probes derived from the two H5N1 strains used in this study, TX37-H5N1 and VN1203-H5N1, were tested for viral receptor binding on the surface of human respiratory organoids. rHAs were precomplexed with anti-Strep-tag II monoclonal antibody (StrepMAB-Classic; IBA Lifesciences, 1.0 μg/ml) and a biotinylated secondary antibody (goat anti-mouse IgG, human adsorbed–biotin, 1:200 dilution) for 30 min at 4°C. After blocking with 10% normal goat serum, tissue sections were incubated for 12–16 h at 4°C with the rHA–antibody complexes. Sections were then washed with PBS and incubated for 30 min at 23–28 °C with streptavidin–HRP (Nichirei Bioscience Inc., Tokyo, Japan). After washing, the sections were developed with DAB-H₂O₂ substrate solution, counterstained with Mayer’s hematoxylin, dehydrated through a graded alcohol series, and cleared in xylene. Histological sections were observed and imaged using a BX43 upright light microscope (Evident Corporation, Tokyo, Japan).

For sialidase treatment, slides were incubated for 16 h at 37°C with neuraminidase from *Vibrio cholerae* (Merck; 1:200 dilution) in 50 mM acetate buffer (pH 5.5). After enzymatic digestion, sections were washed and endogenous peroxidase activity was blocked.

### Cell lines

Madin-Darby canine kidney (MDCK) cells were kindly provided by Dr. Yoshihiro Kawaoka (University of Wisconsin–Madison, USA). MDCK cells were cultured in Eagle’s minimal essential medium (MEM) containing 5% newborn calf serum (NCS). HEK 293S GnTI–/– cells^28^, which lack a functional N-acetyl-glucosaminyltransferase I gene, were maintained in pyruvate-free Dulbecco’s modified Eagle’s medium (Thermo Fisher Scientific) supplemented with 0.3 mg/ml L-glutamine (Nacalai Tesque, Inc., Kyoto, Japan), 100 U/ml penicillin G (Meiji Seika Pharma Co., Ltd., Tokyo, Japan), 0.1 mg/ml streptomycin (Meiji Seika Pharma), 8 µg/ml gentamicin (Takata Pharmaceutical Co., Ltd., Saitama, Japan), and 5% fetal calf serum (Merck KGaA, Darmstadt, Germany). Cells were incubated at 37°C with 5% CO2.

### Human iPS cells

The human induced pluripotent stem (iPS) cell line 1383D6 (provided by Dr. Masato Nakagawa, Kyoto University) was maintained on 0.5 μg/cm^2^ recombinant human laminin 511 E8 fragments (iMatrix-511, Cat. # 892 012, Nippi Inc., Tokyo, Japan) with StemFit AK02N medium (Cat. # RCAK02N, Ajinomoto Healthy Supply Co., Inc., Tokyo, Japan). Cells were passaged every 6 days. For cell passaging, human iPS cell colonies were treated with TrypLE Select Enzyme (Cat. # 12563029, Thermo Fisher Scientific) for 10 min at 37°C and seeded with StemFit AK02N medium containing 5 μM Y-27632 (Cat. # 034-24024, FUJIFILM Wako Pure Chemical Corp., Osaka, Japan).

### Respiratory organoids

To initiate differentiation, human iPS cell colonies were treated with TrypLE Select Enzyme (Thermo Fisher Scientific) for 10 min at 37°C. After centrifugation, cells were seeded onto Matrigel Growth Factor Reduced Basement Membrane (Cat. # 354230, Corning Inc., Corning, NY, USA)-coated cell culture plates (2.0×10^5^ cells/4 cm^2^) and cultured for 2 days. The differentiation of the respiratory organoids was performed in serum-free differentiation (SFD) medium composed of DMEM/F12 (3:1) (Cat. # 044-29765, FUJIFILM Wako Pure Chemical; Cat. # 11320033, Thermo Fisher Scientific) supplemented with N2 (Cat. # 141-08941, FUJIFILM Wako Pure Chemical), B-27 Supplement Minus Vitamin A (Cat. # 12587001, Thermo Fisher Scientific), ascorbic acid (Cat. # ST-72132, STEMCELL Technologies Inc., Vancouver, BC, Canada, 50 μg/mL), 1× GlutaMAX (Cat. # 35050-061, Thermo Fisher Scientific), 1% monothioglycerol (Cat. # 195-15791, FUJIFILM Wako Pure Chemical), 0.05% BSA (Cat. # 820024, Sigma-Aldrich Inc., St. Louis, MO, USA), and 1× penicillin/streptomycin. During days 0-1 of differentiation, the cells were cultured with SFD medium supplemented with 10 μM Y-27632 (Cat. # 034-24024, FUJIFILM Wako Pure Chemical) and 100 ng/mL recombinant Activin A (Cat. # 338-AC-01M, R&D Systems Inc., Minneapolis, MN, USA). During days 1–3 of differentiation, the cells were cultured with SFD medium supplemented with 10 μM Y-27632 (FUJIFILM Wako Pure Chemical), 100 ng/mL recombinant Activin A (Cat. # 338-AC-01M, R&D Systems), and 1% FBS. Between days 3–5 of differentiation, the cells were cultured in SFD medium supplemented with 1.5 μM Dorsomorphin dihydrochloride (Cat. # 047-33763, FUJIFILM Wako Pure Chemical) and 10 μM SB431542 (Cat. # 037-24293, FUJIFILM Wako Pure Chemical) for 24 h, and then SFD medium supplemented with 10 μM SB431542 and 1 μM IWP2 (Cat. # 04-0034, Stemolecule; Reprocell Inc., Yokohama, Japan) for another 24 h. During days 5–12 of differentiation, the cells were cultured with SFD medium supplemented with 3 μM CHIR99021 (Cat. # 034-23103, FUJIFILM Wako Pure Chemical), 10 ng/mL human FGF10 (Cat. # AF-100-26, PeproTech Inc., Cranbury, NJ, USA), 10 ng/mL human FGF7 (Cat. # AF-100-19, PeproTech), 10 ng/mL human BMP4 (Cat. # 120-05ET, PeproTech), 20 ng/mL human EGF (Cat. # AF-100-15, PeproTech), and all-trans retinoic acid (Cat. # R2625 ATRA, Sigma-Aldrich). On day 12 of differentiation, the cells were dissociated and embedded in the Matrigel Growth Factor Reduced Basement Membrane to generate organoids. During days 12– 20 of differentiation, the organoids were cultured in SFD medium containing 3 μM CHIR99021, 10 ng/mL human FGF10, 10 ng/mL human FGF7, 10 ng/mL human BMP4, and 50 nM ATRA. On day 20 of differentiation, the organoids were recovered from the Matrigel, and the resulting organoid suspension (small free-floating clumps) was seeded onto Matrigel-coated cell culture plates. During days 20–30 of differentiation, the organoids were cultured in SFD medium containing 50 nM dexamethasone (Cat. # S1322, Selleck Chemicals LLC, Houston, TX, USA), 0.1 mM 8-bromo-cAMP (Cat. # 1140/50, Tocris Bioscience), and 0.1 mM IBMX (3-isobutyl-1-methylxanthine) (Cat. # 095-03413, FUJIFILM Wako Pure Chemical).

### Immunofluorescence staining

For immunofluorescence staining of respiratory organoids, cells were fixed with 4% paraformaldehyde in PBS at 4°C. Respiratory organoids were harvested and prepared as paraffin sections (approximately 15 μm). Paraffin was removed using xylene and the sections were rehydrated with different percentages of ethanol. Antigen retrieval was performed with 0.1%-tTBS (10×) (pH 7.4) (Cat. # 12750-81, Nacalai Tesque). For blocking non-specific staining, the slides were incubated in Blocking One (Cat. # 03953-066, Nacalai Tesque) for 10 min at room temperature. The primary antibody, mouse anti-acetylated alpha-Tubulin (Cat. # 66200-1-Ig, Proteintech Group Inc., Rosemont, IL, USA), was applied and incubated for 16 h at 4°C. After three washes with 1× PBS (Cat. # 14249-24, Nacalai Tesque), the slides were incubated with Alexa Fluor 488-conjugated anti-mouse IgG (H+L) secondary antibody (Cat. # A21202, Thermo Fisher Scientific) for 45 minutes at room temperature. Slides were then washed three times for 5 min each with PBS. Sections were washed, mounted with ProLong Glass Antifade Mountant with NucBlue Stain (Cat. # P36985, Thermo Fisher Scientific) and DAPI (Cat. # 12593-64, Nacalai Tesque), and analyzed using an inverted laser scanning confocal microscopy system (FV3000, Evident).

### Viruses

TX37-H5N1 was kindly provided by the Centers for Disease Control and Prevention, USA. Cal04-H1N1 was kindly provided by Dr. Yoshihiro Kawaoka (University of Wisconsin–Madison, USA). TX37-H5N1, VN1203-H5N1, and Cal04-H1N1 viruses were propagated in MDCK cells using Eagle’s minimum essential medium (MEM) supplemented with 0.3% BSA and 1 μg/ml N-p-tosyl-L-phenylalanine chloromethyl ketone (TPCK)-treated trypsin. Virus stocks were aliquoted and stored at –80 °C until use. All experiments involving live viruses were conducted in biosafety level 3 (BSL-3) containment facilities and were approved by the Institutional Review Board of the Research Institute for Microbial Diseases, The University of Osaka (protocol number: BIKEN-00311).

### Virus growth kinetics in organoids

Human respiratory organoids were infected in triplicate with virus at a dose of 8 × 10^4^, 8 × 10³, or 8 × 10^2^ PFU. After incubation at 37°C for 1 h, the virus inoculum was replaced with culture medium, followed by further incubation at 37°C with 5% CO₂. Culture supernatants (n = 3) were collected at 0, 6, 12, 24, 48, and 72 h.p.i. Virus titers at the indicated time points were determined by plaque assay in MDCK cells. Data represent the mean ± s.d. (n = 3) from a single experiment.

### Antivirals

The neuraminidase inhibitor oseltamivir acid and the polymerase inhibitor favipiravir (T-705) were purchased from MedChemExpress (Cat. # HY-13318) and Selleck Chemicals (Cat. # S7975), respectively. The endonuclease inhibitor baloxavir acid was obtained from Shionogi & Co., Ltd. (Osaka, Japan). All compounds were dissolved in dimethyl sulfoxide (DMSO; FUJIFILM Wako) for use in experiments.

### Histological examination

On day 3 after virus infection, virus-infected or uninfected organoids were fixed with 4% paraformaldehyde and embedded in paraffin. Influenza virus nucleoprotein (NP) antigens were stained with antiserum from rabbits immunized with an NP synthetic peptide (AFTGNTEGRTSDMR at positions 428–441 of the NP sequence^30^; GenBank accession number, ADC34563) after heat antigen retrieval for 15 min in 0.01 M citrate-phosphate buffer. After incubation with anti-rabbit immunoglobulin antibody conjugated with horseradish peroxidase (Cat. #724142, Nichirei Bioscience Inc.), NP was detected with diaminobenzidine (Cat. #725191, Nichirei Biosciences Inc.).

### Measurement of cytokines in cell supernatants

To measure cytokine concentrations, cell supernatants from virus-infected respiratory organoids were collected and inactivated with β-propiolactone (Cat. # 168-21011, FUJIFILM Wako) prior to analysis. Cytokine levels were then measured using the Human Anti-Virus Response Panel (Cat. # 740349, Biolegend, Inc., San Diego, CA, USA). Flow cytometry was performed according to the manufacturer’s instructions using a MACSQuant Analyzer 10 Flow Cytometer (Miltenyi Biotec GmbH, Bergisch Gladbach, Germany).

### RNA sequencing analysis

Human respiratory organoids were infected with TX37-H5N1, VN1203-H5N1, or Cal04-H1N1 at a dose of 8 × 10^3^ PFU and total RNA was extracted from respiratory organoids (n=3 in each group) at 24, 48, and 72 h.p.i. using an RNeasy Mini Kit (Qiagen, Hilden, Germany), according to the manufacturer’s instructions. All samples were processed and sequenced together in a single batch, ensuring consistency across conditions and time points. For RNA sequencing analysis of virus-infected organoids, library preparation was performed using a TruSeq stranded mRNA Library Prep kit (Illumina Inc., San Diego, CA, USA) according to the manufacturer’s instructions. Sequencing was performed on a NovaSeq X Plus sequencer (Illumina) in a 101-bp single read mode. After adapter trimming by Trimmomatic (v0.39), sequenced reads were mapped to the reference genome sequences (GRCh38) using HISAT2 version 2.1.0. The fragments per kilobase of exons per million mapped fragments (FPKMs) were calculated using Cufflinks version 2.2.1. The raw counts were calculated using featureCounts version 2.0.6. Differential gene expression analysis was performed using DESeq2^31^, with viral strain and replicate included as variables in the design formula (design = ∼ virus_strain + replicate). Differentially expressed genes (DEGs) were defined by an absolute log2(fold change) > 0.5 and an adjusted *p*-value < 0.05. Principal component analysis (PCA) was performed using scikit-learn (v1.0.2) to visualize sample clustering. Inference of pathway activity and transcription factor activity was performed using decoupleR^32^ with the univariate linear model (ULM) method. Pathway activation scores were calculated using Pathway RespOnsive GENes (PROGENy)^33^, while transcription factor (TF) activity was inferred using Collection of Transcription Regulation Interactions (CollecTRI)^34^.

### Single cell RNA sequencing analysis

Human respiratory organoids were infected with TX37-H5N1, VN1203-H5N1, or Cal04-H1N1 at a dose of 8 × 10³ PFU and incubated at 37°C in 5% CO₂. At 24 h.p.i., the organoids were fixed with 4% formaldehyde, and fixed-cell samples were prepared according to the protocol for Fixation of Cells & Nuclei for Chromium Fixed RNA Profiling (10x Genomics, Pleasanton, CA, USA). Single-cell RNA sequencing was performed using the Chromium Fixed RNA Kit, Human Transcriptome, 4rxns x 4 BC and a custom probe set targeting the influenza virus M gene (Supplementary Table 2). Approximately 2,000,000 – 3,000,000 fixed human cells were hybridized for 18 h at 42°C with whole-transcriptome probe pairs. Following post-hybridization washes, the cells were encapsulated into gel beads-in-emulsion (GEMs) using the Chromium X system and Chip Q. Within each GEM, ligated probe pairs were barcoded using 10x GEM Barcodes and Unique Molecular Identifiers (UMIs), followed by heat denaturation and recovery. Barcoded products were pre-amplified and purified using SPRIselect beads. Final libraries were generated by sample index PCR with the Dual Index Kit TS Set A and further purified by size selection. Library quality and size distribution were assessed using an Agilent Bioanalyzer, and library concentrations were determined with the KAPA Library Quantification Kit (Roche, Basel, Switzerland). Libraries were sequenced on an Illumina NovaSeq X Plus platform in paired-end mode (Read 1: 28 bp, Read 2: 90 bp). Demultiplexing, alignment, and gene expression quantification were performed using Cell Ranger v9.0.1 (10x Genomics), aligned to a custom reference genome based on the human genome (GRCh38-2024-A) supplemented with influenza virus sequence. Filtered feature-barcode matrices were generated for downstream transcriptomic analyses.

Samples were aggregated and normalized using the Cell Ranger aggr pipeline. In total, four libraries were prepared, including three from virus-infected samples (TX37-H5N1, VN1203-H5N1, and Cal04-H1N1) and one from mock-infected control, and sequenced in a single batch to minimize technical variation. Quality control was performed as previously described^35^. Briefly, cells with low gene counts, low total unique molecular identifier (UMI) counts, or high mitochondrial content were identified as outliers using 5 median absolute deviation (MAD)-based cut-offs. Cells exceeding these cut-offs were excluded from subsequent analyses. In addition, cells with a mitochondrial count percentage exceeding 8% were excluded. Genes expressed in fewer than 20 cells were also excluded. Doublet detection was performed using the scDblFinder^36^. Dimensionality reduction and visualization were performed using UMAP. Cell-type annotation was performed using CellTypist^37,38^ with cell types from human embryonic and fetal lungs^12^. Cells were classified as infected if influenza virus transcript levels (detected using a probe for the viral M gene; see Supplementary Table 2) exceeded 0.05% or 0.01% of total host mRNA per cell. Transcription factor activity at the single cell level was inferred using CollecTRI^34^ and decoupleR^32^ with the univariate linear model (ULM) approach.

### Statistical analyses

All data are presented as means ± standard deviation (s.d.). Statistical analyses were performed using GraphPad Prism version 9.5.1. Virus growth kinetics in organoids were compared by using a two-way ANOVA followed by Dunnett’s multiple comparisons test. Cytokine quantification and virus titers under antiviral treatment were compared by using a one-way ANOVA followed by Tukey’s multiple comparisons test. A *P* value of <0.05 was considered statistically significant.

## Data Availability

The raw sequencing data generated in this study have been deposited in the DDBJ Sequence Read Archive (DRA) under Bioproject: PRJDB37933 (PSUB043713).

## Acknowledgments

We thank Susan Watson for scientific editing. We also thank Ayaka Sakamoto (Institute of Science Tokyo) and Naoko Yasuhara (Kyoto University) for generating and analysing respiratory organoids, Itsuki Anzai, Tomomi Kirino, Hina Miyazaki, Sara Yoshimoto, Nicholas Yamahoki, and Shota Morimoto, and the NGS Core laboratory, Genome Information Research Center, Research Institute for Microbial Diseases, The University of Osaka, for technical assistance. In addition, we thank Yoshihiro Kawaoka (University of Wisconsin–Madison) for kindly providing MDCK cells and Cal04-H1N1 virus, the Centers for Disease Control and Prevention for kindly providing TX37-H5N1 virus. Figure 2A was created with BioRender (https://biorender.com), and Figures 6A and 7A were obtained from TogoTV (© 2016 DBCLS TogoTV, CC-BY-4.0). This study was supported by JSPS KAKENHI Grants-in-Aid for Scientific Research (C) [JP24K09264 (to S.S.)]; the Japan Agency for Medical Research and Development (AMED) under Grant Numbers JP223fa627005 (to Y.S.), JP243fa627003h0003 and JP24gm1810009 (to E.K.), JP223fa627002 and JP233fa827018 (to T.W.); AMED Advanced Research and Development Programs for Medical Innovation (AMED-CREST) [JP21gm1610005 (to K.T.) and JP22gm1610010 (to T.W.)]; the Science and Technology Agency (JST) Moonshot R&D grant (JPMJMS2025 to E.K.); and the Takeda Science Foundation (to T.W.).

## Contributions

Author contributions are provided according to Contributor Roles Taxonomy (CRediT). Conceptualization: S.S. and T.W. Data curation: H.S., D.M., and E.K. Formal analysis: H.S., D.M., and E.K. Funding acquisition: S.S., Y.S., E.K., K.T., and T.W. Investigation: S.S., M.Y., R.H., T.M., D.K., T.H., and D.M. Methodology: S.S., T.M., T.H., M.S., and K.T. Project administration: S.S. and T.W. Resources: N.I., M.T.Q.L., A.T., Y.S., K.T., and T.W. Software: H.S. and E.K. Supervision: E.K., K.T., and T.W. Validation: S.S., H.S., M.Y., R.H., T.M., and T.H. Visualization: S.S., H.S., and M.Y. Writing—original draft: S.S., H.S., M.Y. R.H., T.M., T.H., and D.M. Writing—review and editing: M.S., N.I., Y.S., E.K., K.T., and T.W. Author contributions to specific experiments: Receptor-binding studies were performed by S.S., M.Y., R.H., T.M., D.K., and T.H. Virus infection and cytokine detection experiments were performed by S.S., M.Y., R.H., M.S., and K.T. Antiviral susceptibility testing experiments were performed by S.S. Bulk RNA-seq and single cell RNA-seq analysis was performed by S.S., H.S., D.M., and E.K.

