## Supplementary Table 2 for "A cattle-derived human H5N1 isolate suppresses innate immunity despite efficient replication in human respiratory organoids"

**Supplementary Table 2 | Probe sequences for influenza virus genome detection.**

| Left Hand Side (LHS) probe | Sequence (5′→3′) |
|---|---|
| IFV-M-LHS | CCTTGGCACCCGAGAATTCCAGACAAAGCGTCTACGCTGCAGTCCT |
| Right Hand Side (RHS) probes | Sequence (5′→3′) |
| IFV-M-RHS_BC001 | 5′Phos-CGCTCACTGGGCACGGTGAGCGTGAACGCGGTTAGCACGTANNACTTTAGGCGGTCCTAGCAA |
| IFV-M- RHS_BC002 | 5′Phos-CGCTCACTGGGCACGGTGAGCGTGAACGCGGTTAGCACGTANNAACGGGAACGGTCCTAGCAA |
| IFV-M- RHS_BC003 | 5′Phos-CGCTCACTGGGCACGGTGAGCGTGAACGCGGTTAGCACGTANNAGTAGGCTCGGTCCTAGCAA |
| IFV-M- RHS_BC004 | 5′Phos-CGCTCACTGGGCACGGTGAGCGTGAACGCGGTTAGCACGTANNATGTTGACCGGTCCTAGCAA |

Custom probe sets were designed to detect transcripts of the influenza virus M gene and were incorporated into the 10x Genomics Fixed RNA Profiling workflow to identify infected cells in human respiratory organoids.
